# Loss of Tubulin Tyrosination in Purkinje Neurons Does Not Cause Their Degeneration

**DOI:** 10.64898/2026.07.21.739800

**Authors:** Shreyangi Chakraborty, Sloane Paulcan, Carsten Janke, Maria M. Magiera

## Abstract

Posttranslational modifications (PTMs) of tubulin have been suggested to form a ‘tubulin code’ to coordinate microtubule functions. While at the molecular level PTMs were demonstrated to act in a target-specific manner, an outstanding question is the selectivity with which they control cellular and physiological functions. For instance, several tubulin PTMs have been linked to neurodegeneration, suggesting that whichever PTM is dysregulated could be sufficient to induce microtubule perturbation and cause neuronal death. Here we test whether perturbation of tyrosination can cause degeneration of Purkinje neurons similar to what happens when the PTM polyglutamylation is abnormally accumulated. Strikingly, this is not the case: deletion of tubulin tyrosine ligase in Purkinje cells, while resulting in loss of tyrosinated tubulin in these neurons, does not affect their survival nor function for more than one year. Our results thus reveal a highly specific role of polyglutamylation in neuronal homeostasis that is not replicated by other tubulin PTMs, suggesting stringent functional specialisation of those PTMs at the physiological level.

## Introduction

Neurons are long-lived, architecturally complex cells that strongly rely on the microtubule cytoskeleton for maintaining their homeostasis and functions. Consequently, aberrant regulation of microtubules or their associated factors lead to perturbation of neuronal functions which often result in neurodegeneration (Penazzi et al, 2016; Sleigh et al, 2019). One of the mechanisms controlling microtubule functions in neurons are posttranslational modifications (PTMs) of tubulins, the microtubule building blocks (Teoh & Bartolini, 2025). On the molecular level, tubulin PTMs can regulate intrinsic microtubule properties (Chen & Roll-Mecak, 2023), or modulate the events taking place on microtubules such as binding of interacting partners (Genova et al, 2023; Krishnan et al, 2026), cargo transport (Bodakuntla et al, 2020; Genova et al, 2023; Gilmore-Hall et al, 2019), or microtubule severing (Lacroix et al, 2010; Szczesna et al, 2022; Valenstein & Roll-Mecak, 2016). The importance of microtubule-based processes for neuronal functions together with the particularly strong enrichment of tubulin PTMs such as detyrosination, acetylation and polyglutamylation on neuronal microtubules have for long suggested that disruption of tubulin PTMs could have a deleterious impact on neuronal functions and eventually cause neurodegeneration. Past work has provided evidence that several tubulin PTMs can, if deregulated, cause neurodegeneration (Pero et al, 2023; Teoh & Bartolini, 2025), however it is not clear whether deregulation of any tubulin PTM can cause neurodegeneration in similar physiological contexts.

The so-far most direct evidence for a perturbed tubulin PTM directly causing neurodegeneration has been provided for polyglutamylation (Bodakuntla et al, 2021a), a major tubulin PTM in the nervous system (Audebert et al, 1993; Audebert et al, 1994; Eddé et al, 1990). Polyglutamylation is strongly increased above wild-type levels in several regions of the nervous system in mice carrying a mutation in the deglutamylase *CCP1* (Bodakuntla et al, 2021b; Magiera et al, 2018; Rogowski et al, 2010). This ‘hyperglutamylation’ leads to a massive and synchronous degeneration of the cerebellar Purkinje cells between three and five weeks after birth, which is why they were originally named Purkinje-Cell Degeneration (*pcd*) mice (Mullen et al, 1976). We have shown that the degeneration can be entirely avoided when the main polyglutamylase in brain, Ttll1 (Janke et al, 2005), is concomitantly inactivated (Bodakuntla et al, 2021b; Magiera et al, 2018). Hyperglutamylation-related degeneration is not restricted to Purkinje cells: other neurons such as mitral cells in the olfactory bulb, motor neurons and other neuronal types (Delis et al, 2004; Greer & Halasz, 1987; Magiera et al, 2018) also degenerate in *pcd* mice. Moreover, we found that in brain regions that are not affected in *pcd* mice, polyglutamylation levels are maintained at near-wild-type levels by a second deglutamylase, CCP6. Accordingly, double-knockout mice lacking both CCP1 and CCP6 (*Ccp1^-/-^/Ccp6^-/-^*) showed more severe neurodegeneration including in those regions (Magiera et al, 2018). This demonstrated that hyperglutamylation is a general cause of neuronal dysfunction and degeneration.

Among all degenerative processes observed in either *Ccp1^-/-^*or *Ccp1^-/-^/Ccp6^-/-^* mice, the rapid degeneration of the Purkinje cells is the most striking and outstanding effect of hyperglutamylation. It highlights the particular susceptibility of these neurons to perturbations of the microtubule cytoskeleton, which also degenerate if other regulators of microtubules are perturbed, such as the neuronal microtubule-associated protein Map1A (Liu et al, 2015). While the detailed molecular and cellular pathways leading to Purkinje-Cell degeneration caused by hyperglutamylation have not yet been unveiled, it has been demonstrated that the transport of multiple cellular cargoes is perturbed in affected neurons (Bodakuntla et al, 2020; Bodakuntla et al, 2021b; Gilmore-Hall et al, 2019; Magiera et al, 2018). At the molecular level, microtubule interactors with key functions in neurons such as the microtubule-associated proteins Map1B, Map2 or Tau (Genova et al, 2023; Krishnan et al, 2026), as well as the microtubule-severing enzyme spastin (Lacroix et al, 2010; Valenstein & Roll-Mecak, 2016) are regulated by polyglutamylation. While genetic experiments already excluded spastin-mediated microtubule severing as cause of the degeneration of Purkinje cells in *Ccp1^-/-^* mice (Magiera et al, 2018), given the multitude of MAPs regulated by polyglutamylation (Krishnan et al, 2026), perturbed binding of one or several neuronal MAPs might be involved in the pathology.

If perturbations of multiple MAPs are part of the molecular mechanism underlying hyperglutamylation-related Purkinje-Cell degeneration, other mechanisms affecting MAP-microtubule interactions might have similar impacts when perturbed. Among the so-far analysed tubulin PTMs, loss of tyrosination has been shown to massively impact the binding of several MAPs to microtubules (reviewed in Sanyal et al, 2023). Moreover, while polyglutamylation often only gradually changes MAP affinities to microtubules, loss of tyrosination can change microtubule-MAP interactions more severely (Bieling et al, 2008; Honnappa et al, 2006; Krishnan et al, 2026; Peris et al, 2006). One would thus expect that perturbations of tyrosination could, similar to polyglutamylation, lead to neurodegeneration, or have an even more striking physiological impact.

*TTL* knockout (*Ttl*^-/-^) mice, which lack this unique enzyme for re-tyrosination of detyrosinated tubulin (Fig. 2A), accumulate detyrosinated and Δ2-tubulin to an extent that tyrosinated tubulin can merely be detected in neurons. *Ttl*^-/-^ mice exhibit massive developmental defects in the brain and die perinatally. Primary neurons that can be obtained from *Ttl*^-/-^ embryos show increased neurite outgrowth and premature differentiation (Erck et al, 2005). The perinatal lethality of *Ttl*^-/-^ mice had so far limited studies on the role of tubulin detyrosination in postnatal brain homeostasis and neurodegeneration. A first study using a conditional *Ttl*-flox mouse to delete TTL postnatally in the hippocampus demonstrated reduced synaptic plasticity in *Ttl^flox/flox^* CaMKIIα-cre^+/-^ mice. Interestingly however, no signs of neurodegeneration were found (Hosseini et al, 2022). The absence of neurodegenerative phenotypes in this mouse model is puzzling, as tyrosination has a strong effect on the interactions of several MAPs with microtubules (Krishnan et al, 2026). Moreover, in Alzheimer’s disease (AD) patient samples, reduced levels of TTL protein have been detected, and loss of TTL function was shown to exacerbate amyloid-β-induced synaptic damage (Peris et al, 2022), which together suggested that decreased tyrosination could be linked to neurodegeneration.

To investigate whether the loss of tubulin tyrosination has the potential to cause neurodegeneration, we knocked out *TTL* in the neurons that have most sensitively reacted to perturbation of polyglutamylation, the Purkinje cells, by using L7-cre (Rico et al, 2004), and investigated Purkinje-cell survival to up to 13 months. Strikingly, while tyrosination was undetectable in Purkinje cells of *Ttl^flox/flox^* L7-cre mice, no degeneration, nor signs of ataxia caused by Purkinje-cell dysfunction were observed. These results show that in contrast to increased polyglutamylation, massive loss of tyrosination does not impede the survival and function of cerebellar Purkinje neurons. Our findings illustrate that tubulin PTMs can fulfil unique physiological roles, and consequently their perturbations lead highly specific pathologies.

## Results

### Loss of TTL does not affect Purkinje-cell survival

To specifically knock out *TTL* in Purkinje cells, we used the L7-cre mouse model that had previously been shown to express cre recombinase exclusively in these cells in the cerebellum (Rico et al, 2004) (Fig. 1A). *Ttl^flox/flox^* L7-cre^+/-^ (referred to as *Ttl^flox/flox^* L7-cre in the manuscript) mice and their *Ttl^flox/flox^*L7-cre^-/-^ control littermates (from now-on referred to as *Ttl^flox/flox^*, or control) were generated by breeding *Ttl^flox/flox^* L7-cre^+/-^ and *Ttl^flox/flox^* animals. This breeding scheme yielded 46.5% *Ttl^flox/flox^* L7-cre animals (over 34 separate litters), close to the expected 50%. *Ttl^flox/flox^* L7-cre mice develop normally, demonstrating that unlike the systemic knockout of *TTL* (Erck et al, 2005), its targeted deletion in Purkinje cells is not lethal.

**Figure 1.**
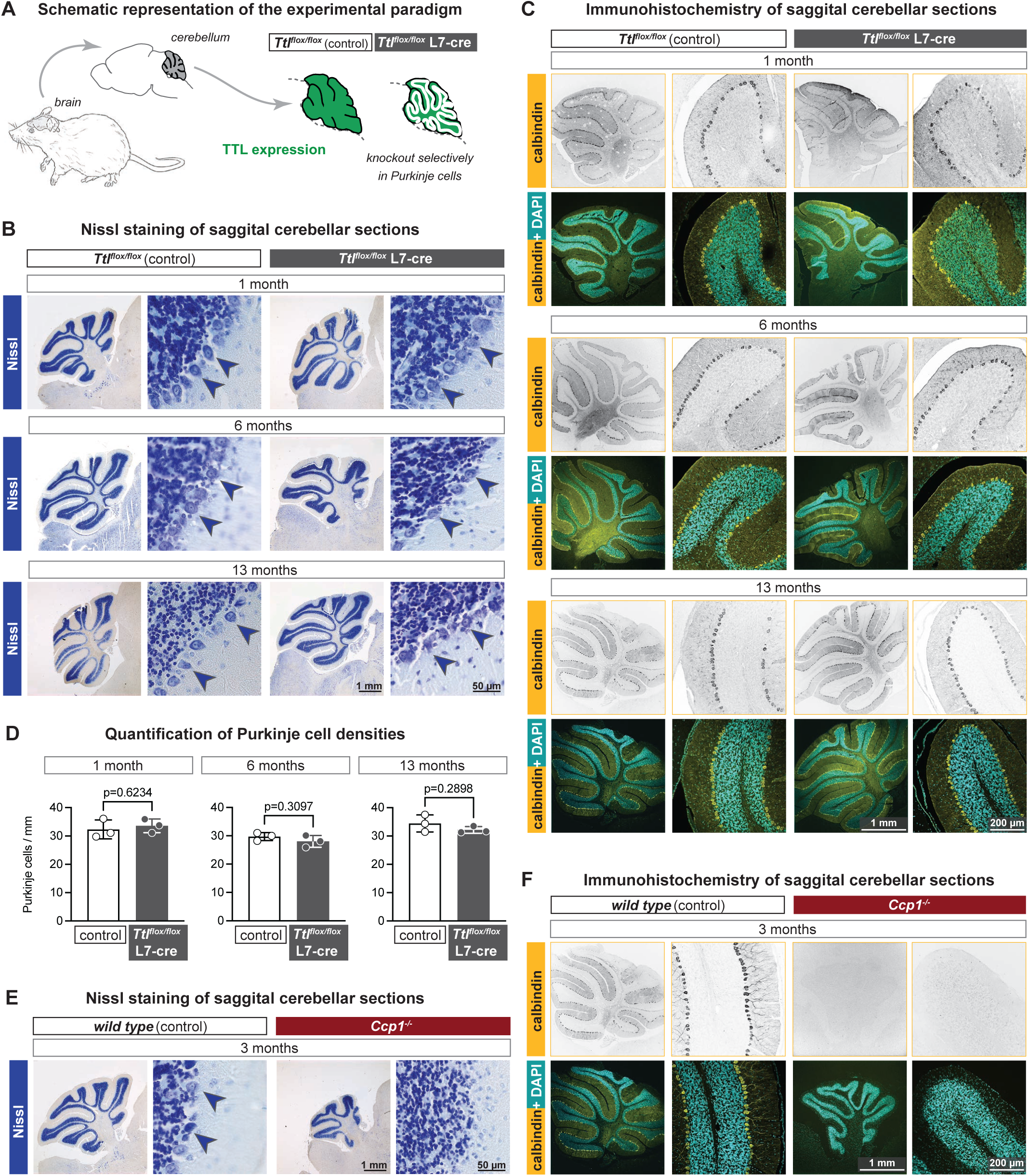
Purkinje cells do not degenerate in ageing *Ttl^flox/flox^* L7-cre mice. **A.** Schematic representation of the experimental paradigm. Tubulin tyrosine ligase (TTL; green) is specifically depleted in Purkinje cells of *Ttl^flox/flox^* mice expressing L7-cre. **B.** Nissel staining of cerebellar sections of control *Ttl^flox/flox^* and *Ttl^flox/flox^*L7-cre mice at 1, 6 and 13 months. Representative Purkinje neurons are highlighted with blue arrows. **C.** Calbindin staining of cerebellar sections of control *Ttl^flox/flox^* and *Ttl^flox/flox^*L7-cre mice at 1, 6 and 13 months. **D.** Quantitation of Purkinje-cell densities in control and *Ttl^flox/flox^*L7-cre mice at 1, 6 and 13 months. Number of Purkinje-cell bodies per length was quantified in two folia per mouse brain. Mean values are determined from three mice. Bars indicate standard deviation. Statistical significance was tested using the unpaired t-test. **E.** Nissel staining of cerebellar sections of control and *Ccp1^-/-^* mice at 3 months. **F.** Calbindin staining of cerebellar sections of control and *Ccp1^-/-^* mice at 3 months. Scale bars: B and E: 1 mm, 50 µm, C and F: 1 mm, 200 µm. 20

In a similar mouse model where the deglutamylase CCP1 was specifically inactivated in Purkinje cells using L7-cre, most Purkinje cells degenerated before 1 month postnatally, similar to the timeline observed in *Ccp1^-/-^* mice (Magiera et al, 2018). We thus argued that if perturbed tyrosination has the potential to cause neurodegeneration, a Purkinje-cell-specific knockout of *TTL* should be sufficient to induce a visible loss of these neurons. However, as the time course of degeneration could be different – for instance much longer - than the rapid degeneration observed in *pcd*/*Ccp1^-/-^* mice (Magiera et al, 2018; Mullen et al, 1976), we analysed *Ttl^flox/flox^* L7-cre mice at 1, 6 and 13 months.

We first examined sagittal cerebellar sections by Nissl staining, which allows to visualise the characteristically large cell bodies of Purkinje neurons (Kreutzberg, 1984). Nissl staining revealed the presence of a seemingly unperturbed Purkinje-cell layer in cerebella of *Ttl^flox/flox^* L7-cre mice, similar to what was observed in *Ttl^flox/flox^*control mice, at all ages investigated (Fig. 1B). We next confirmed the presence of Purkinje cells with anti-calbindin antibody, a marker specific of Purkinje neurons (Sequier et al, 1988) (Fig. 1C). Calbindin staining confirmed the presence of a morphologically normal Purkinje-cell layer in 1-, 6- and 13-month-old *Ttl^flox/flox^* L7-cre mice. Using calbindin staining, we quantified the number of Purkinje cells per length within two folia of the Purkinje-cell layer, revealing that Purkinje neuron density is virtually unchanged between control and *Ttl^flox/flox^* L7-cre mice at all ages (Fig. 1D).

Our findings suggest that Purkinje cells develop normally and do not degenerate in aged mice lacking tyrosinated tubulin in Purkinje cells. This stands in stark contrast to the *Ccp1^-/-^* mice which completely lack Purkinje cells at 3 months, as visualised with the Nissl staining (Fig. 1E) and calbindin immunostaining (Fig. 1F) of *Ccp1^-/-^* cerebella.

### TTL deletion specifically reduces levels of tyrosinated tubulin in Purkinje neuron

As previously demonstrated in *Ttl^-/-^*mice, loss of TTL leads to an almost complete absence of tyrosinated tubulin (Erck et al, 2005), which is induced by the unopposed activity of detyrosinating enzymes (Fig. 2A). By contrast, our selective knockout of *TTL* only in Purkinje cells (Fig. 1A; 2B) made it impossible to characterise the resulting decrease in tyrosination by immunoblot. We thus visualised the PTM status of Purkinje cells on cerebellar sections by immunohistochemistry using well-characterised PTM-specific antibodies.

**Figure 2.**
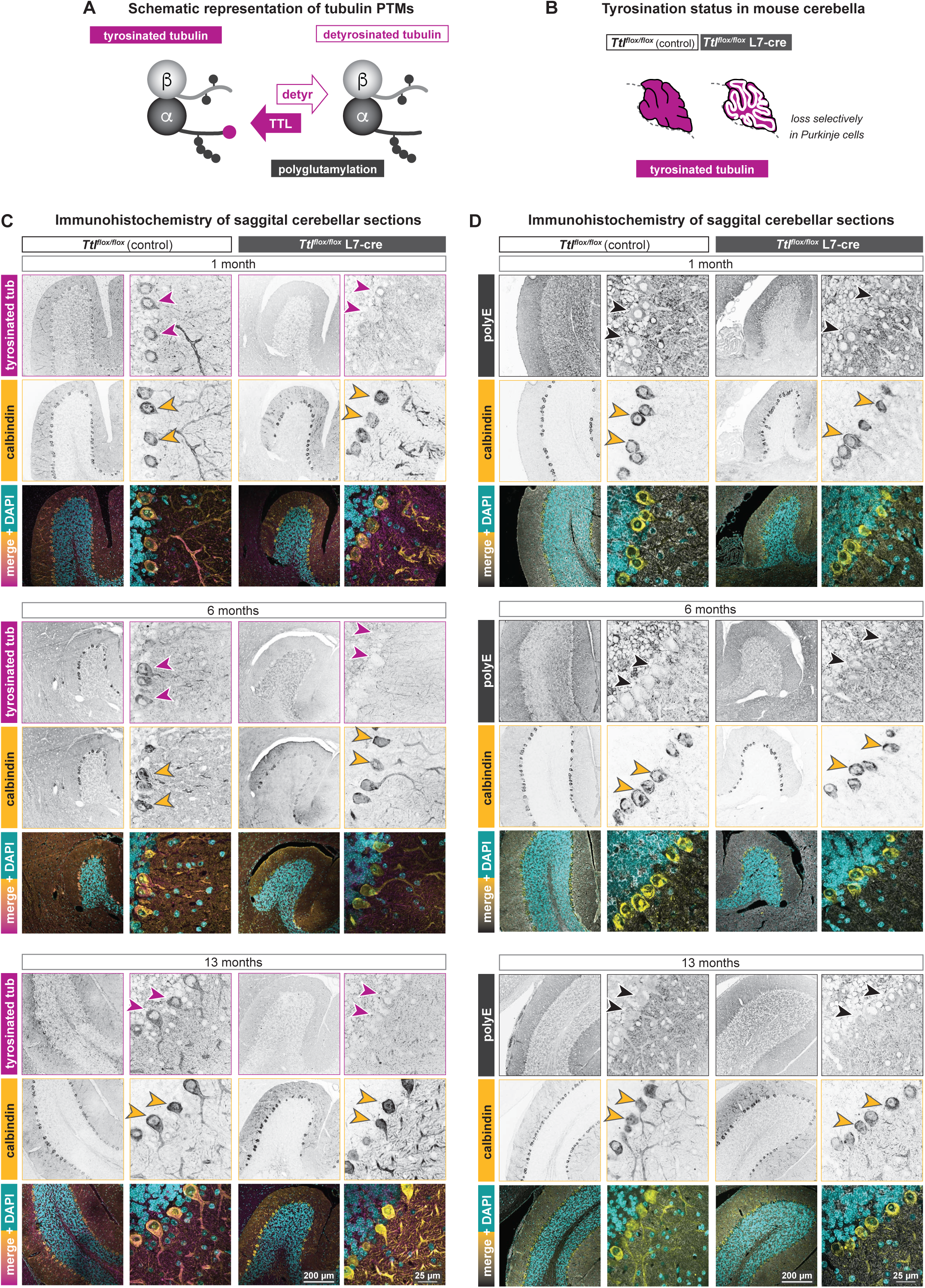
Purkinje cells in *Ttl^flox/flox^* L7-cre mice lack tyrosinated tubulin. **A.** Schematic representation of tubulin PTMs affecting the α-tubulin carboxy-terminal tail in neurons, detyrosination/tyrosination (magenta). Loss of TTL leads to a shift of the balance toward detyrosinated tubulin. Polyglutamylation (black) is not expected to be affected. **B.** Schematic representation of how the selective deletion of TTL (Fig. 1A) would lead to a selective loss of tyrosination in Purkinje cells of *Ttl^flox/flox^* mice expressing L7-cre (magenta). **C.** Tyrosination and calbindin staining of cerebellar sections of control *Ttl^flox/flox^*and *Ttl^flox/flox^* L7-cre mice at 1, 6 and 13 months. Calbindin highlights Purkinje cells (yellow arrows), which show a clear signal for tyrosinated tubulin in controls, while no signal was detected in L7-cre brains at all ages (purple arrows). **D.** Polyglutamylation and calbindin staining of cerebellar sections of control *Ttl^flox/flox^* and *Ttl^flox/flox^* L7-cre mice at 1, 6 and 13 months. Polyglutamylation in Purkinje cells (yellow arrows) with no differences between genotypes at all ages (black arrows).

Immunostaining of sagittal cerebellar sections of 1-, 6- and 13-month-old animals with the TUB-1A2 antibody (Banerjee et al, 2010; Kreis, 1987), showed a strong and specific decrease of tyrosination levels in Purkinje neurons of *Ttl^flox/flox^* L7-cre mice at all three ages as compared to controls (Fig. 2C, purple arrows). These results confirm that the selective knockout of *TTL* in Purkinje cells leads to the selective loss of tyrosinated tubulin specifically in these neurons. *Ttl^flox/flox^* L7-cre mice therefore experience a sustained reduction of tubulin tyrosination throughout life.

Because polyglutamylation occurs in the proximity of detyrosination on the carboxy-terminal tails of α-tubulin (Fig. 2A), and a previous report showed an interplay between these two PTMs (Ebberink et al, 2023), we wanted to verify whether the observed changes in the levels of tyrosinated tubulin in the Purkinje cells of *Ttl^flox/flox^* L7-cre mice also affect polyglutamylation. Polyglutamylation was assessed with the polyE antibody, which detects polyglutamate side chains longer than three glutamates (Rogowski et al, 2010; Shang et al, 2002). PolyE staining showed no difference between *Ttl^flox/flox^* L7-cre and *Ttl^flox/flox^* Purkinje neurons at all ages (Fig. 2D, black arrowheads), indicating that polyglutamylation levels did not change detectibly despite the massive loss of tyrosination.

Collectively, the immunohistological characterisation of tubulin PTMs in Purkinje cells demonstrates that L7cre-mediated knockout of *TTL* in these neurons leads to a massive loss of tubulin tyrosination. The change in this PTM is specific, as polyglutamylation was not altered in Purkinje neurons of these mice.

### TTL depletion in Purkinje cells does not lead to ataxia in mice

In *pcd* (or *Ccp1^-/-^*) mice, a characteristic early occurrence of an ataxic phenotype clearly correlates with the early degeneration of Purkinje cells (Mullen et al, 1976), which are the sole output of the cerebellum. Moreover, the mere dysfunction of these neurons without full degeneration can also lead to ataxia (Jaarsma et al, 2024). To test whether Purkinje neurons in *Ttl^flox/flox^* L7-cre mice are functional, we assessed the motor performance of 16-month-old *Ttl^flox/flox^*L7-cre mice and their control littermates. Both the hind limb clasping (Fig. 3A) and the observation of the mice when freely moving in their cages (Suppl. Movie 1) showed no signs of motoric dysfunction of *Ttl^flox/flox^* L7-cre mice, while *Ccp1^-/-^* mice are strongly affected.

**Figure 3.**
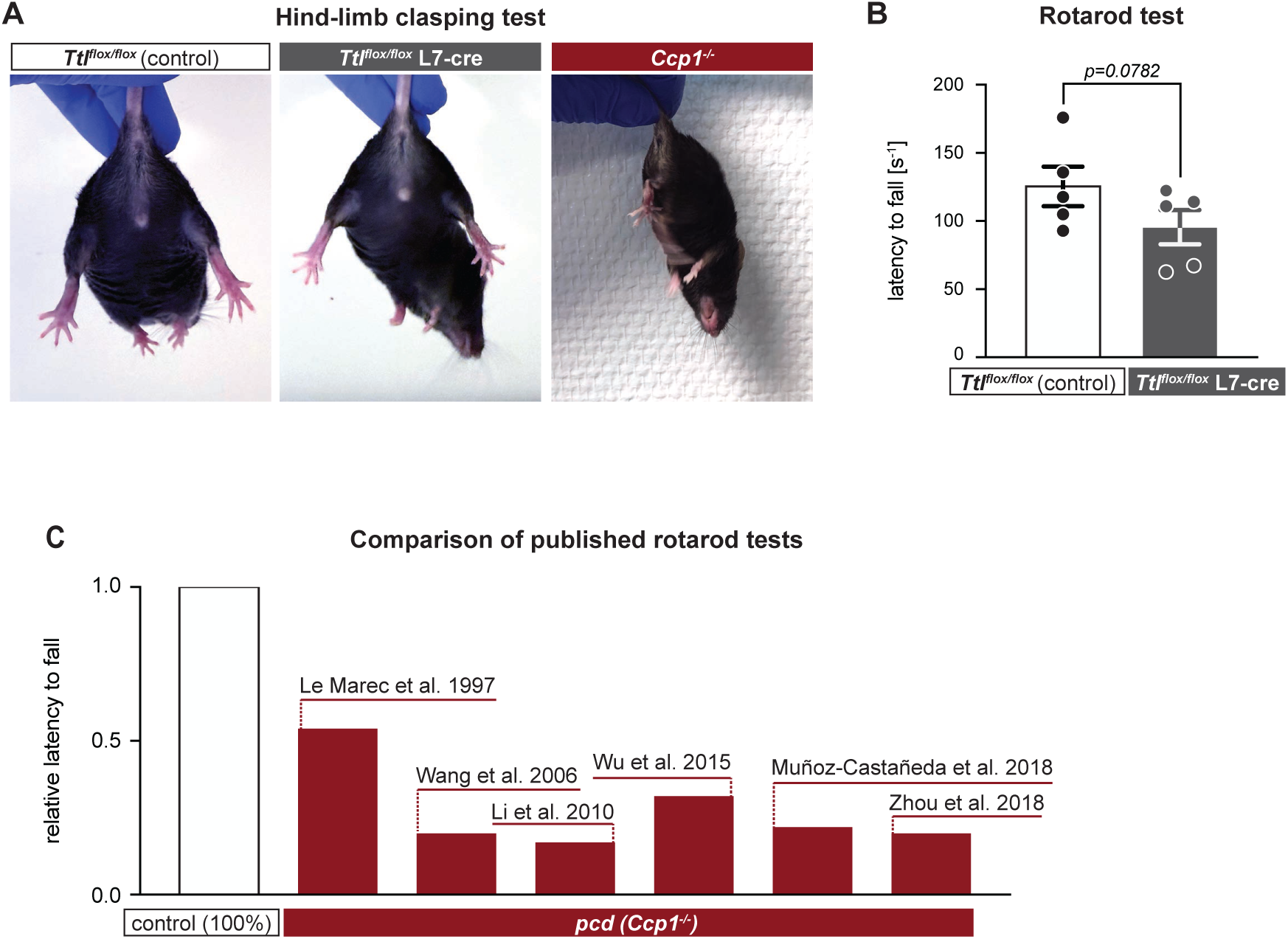
Ttl^flox/flox^ L7-cre mice do not exhibit ataxic phenotypes. **A.** Photographs of control (*Ttl^flox/flox^*), *Ttl^flox/flox^* L7-cre and *Ccp1^-/-^* mice held by the tail. Note that *Ttl^flox/flox^* L7-cre animal, unlike the *Ccp1^-/-^* mouse, does not show hind limb clasping. **B.** Quantification of the latency to fall during a rotarod test of 16-month-old control (*Ttl^flox/flox^*) and *Ttl^flox/flox^* L7-cre animals. The bar graph represents mean of five mice tested, whiskers are SEM, points are individual animals tested. Statistical significance was tested using the unpaired t test. **C.** Compilation of published results of rotarod tests evaluating the latency to fall of *Ccp1 ^/-^* (or *pcd*) mice. For comparability, relative values were used (controls of all studies were set to 1). Animals of the following ages were tested in the respective studies: 2-month-old (Le Marec & Lalonde, 1997), 3-month-old (Wang et al, 2006), 40-day-old (Li et al, 2010), 2- month-old (Wu et al, 2015), 1-month-old (Muñoz-Castañeda et al, 2018), 1-month-old (Zhou et al, 2018).

Ataxia can be measured more quantitatively by the time animals can remain on a rotating rod in the so-called rotarod test. We used this to test whether we can detect a change in motor performance of *Ttl^flox/flox^*L7-cre mice. 16-month-old *Ttl^flox/flox^* L7-cre mice showed a slight, though not significant reduction of time spent on the rotating bar and compared to control littermates, suggesting the absence of ataxia (Fig. 3B). This stands in stark contrast to the massive ataxic phenotypes observed in *pcd* mice in past studies, which were consistently performed with much younger animals (2-3 months) (Le Marec & Lalonde, 1997; Wang et al, 2006; Wu et al, 2015), or even as young as 1 month (Li et al, 2010; Muñoz-Castañeda et al, 2018; Zhou et al, 2018) (Fig. 3C). Our rotarod data thus show that *Ttl^flox/flox^* L7-cre mice maintain normal motor behaviour throughout adulthood, suggesting that Purkinje cells remain functional despite the absence of tyrosinated α-tubulin in these neurons.

## Discussion

Most α-tubulins in mammals carry a gene-encoded carboxy-terminal tyrosine, which can be enzymatically removed by several enzymes known as detyrosinases (Aillaud et al, 2017; Landskron et al, 2022; Nicot et al, 2023; Nieuwenhuis et al, 2017). However, the enzymatic re-addition of tyrosine is catalysed by only one single enzyme: tubulin-tyrosine ligase or TTL (Ersfeld et al, 1993). Deletion of this unique enzyme causes a massive loss of tyrosinated tubulin in most cells of the organism, and there is almost no detectable tyrosinated tubulin present in neurons of *Ttl^-/-^* mice (Erck et al, 2005). The tyrosinated form of α-tubulin has so far been recognised as important molecular requisite of microtubule interactions with several associated proteins. For instance, tyrosination is essential for the recruitment of CAP-Gly proteins (Honnappa et al, 2006), thus playing key roles in the correct intracellular functioning of proteins such as CLIP170 and p150^glued^ (Bieling et al, 2008; Peris et al, 2006), or in controlling the activity of microtubule depolymerases from the kinesin-14 family (Peris et al, 2009). Moreover, when testing a large panel of MAPs for their sensitivity to PTMs on the microtubules they interact with, tyrosination showed strong effects on microtubule binding of several MAPs, while the regulatory effect of polyglutamylation was less pronounced (Krishnan et al, 2026).

In the light of the strong impact of α-tubulin tyrosination status on key molecular processes, we here investigated the importance of this tubulin PTM in neurons, and its potential to cause neurodegeneration. Following our previous work where we had demonstrated that perturbed polyglutamylation causes massive neurodegeneration of the cerebellar Purkinje cells (Magiera et al, 2018), we here generated a similar mouse model in which the knockout of *TTL* is selectively induced in Purkinje neurons. Despite the almost complete loss of tyrosinated tubulin, Purkinje cells showed no sign of degeneration even after more than one year. This tolerance to the massive perturbation of this PTM stands in stark contrast to the deleterious effect the absence of TTL has on brain development (Erck et al, 2005), or to the neurodegeneration caused by loss of CCP1 (Magiera et al, 2018; Shashi et al, 2018).

Our findings underscore that at the physiological level, tubulin PTMs have context-dependent roles. This is most likely based on the specific molecular processes they control (Krishnan et al, 2026). One could hypothesise that loss of tyrosination, which is deleterious for processes that require microtubule dynamics regulated by CAP-Gly proteins such as Clip170 or depolymerising kinesins (Bieling et al, 2008; Peris et al, 2006), is less problematic for differentiated neurons with more stable microtubule tracks that support cargo transport. It might even favour these processes, given initial observations showing that detyrosinated microtubules preferentially interact with kinesin-1, a key molecular motor in axonal transport (Sirajuddin et al, 2014).

Importantly, we observed here and in our previous work on *Ccp1^-/-^* mice (Magiera et al, 2018) that other tubulin PTMs do not vary when a single PTM is perturbed. This underpins that PTMs appear to not compensate for each other at longer time scales, while a short-term interplay between tyrosination and glutamylation has been observed *in vitro* and in cells after overexpression of polyglutamylating enzymes (Ebberink et al, 2023).

While our study establishes upregulated polyglutamylation as a so-far unique PTM causing neurodegeneration, it does not exclude the possibility that other tubulin PTMs play modulatory roles in neuropathies. In Alzheimer’s disease for instance, reduced expression of *TTL* and reduced levels of tyrosinated tubulin correlated with amyloid-β-induced synaptic damage, suggesting that tyrosination may protect against neurodegeneration in specific contexts (Peris et al, 2022). Also, increased acetylation in Alzheimer’s disease brains was suggested to compensate for microtubule loss by potentially stabilising microtubules to help maintaining cargo transport (Hempen & Brion, 1996).

In conclusion, we demonstrate that the almost complete loss of tubulin tyrosination in Purkinje neurons does not induce their degeneration, while it had been shown to have a deleterious effect on neurodevelopment (Erck et al, 2005). Increased polyglutamylation, by contrast, allows the nervous system to develop rather normally but leads to massive degeneration of differentiated neurons (Bodakuntla et al, 2021a). Our work thus emphasises that tubulin PTMs have, similar to the unique effects they have at the molecular scale (Krishnan et al, 2026), clearly distinguishable and functionally independent physiological roles. Perturbations in PTMs have thus unique pathological outcomes, even if they happen in the same cell types. We thus provide evidence for exclusive physiological roles of tubulin PTMs, pointing to the possibility to target specific tubulin PTMs as a means to selectively treat pathologies.

## Acknowledgments

This work was supported by the Institut Curie, the Centre National de la Recherche Scientifique, the European Research Council (ERC-2022-SYG) under the European Union’s Horizon 2020 research and innovation programme (grant agreement N°101071583 ‘TUBULINCODE’), the French National Research Agency (ANR) award ANR-20-CE13-0011, the Fondation pour la Recherche Medicale (FRM) grants MND202003011485 and EQU202203014694. MMM was supported by the Fondation Vaincre Alzheimer FR-16055p and the France Alzheimer grant 2023. SC was supported by a PSL-Biogen PhD grant via a CIFRE contract.

We thank V. Dangles-Marie, C. Alberti, E. Belloir, C. Jouhanneau, K. Belloul, A. Luber and L. Lebrun for help with animal generation; F. Pouzoulet from the RadeXp platform; C-Y. Chang for help with the optimisation of the immunohistochemistry protocol; L. Besse and M.-N. Soler from the imaging platform PICT-IBiSA@Orsay for help with imaging and image analysis; and S. Leboucher and A. Starck from histology platform, all from the Institut Curie, Orsay, for technical assistance.

## Material and Methods

### Mouse lines

Animal care and use were carried out in accordance with the recommendations of the European Community (2010/63/UE). Experimental procedures were specifically approved by the ethics committee of the Institut Curie CEEA-IC #118 (authorisations #04395.03 and APAFIS #37315-2022051117455434 v2 given by the National Authority) in compliance with the international guidelines.

*Ttl^flox/flox^* mice (Bradley et al, 2012; Skarnes et al, 2011) were generated and distributed by The Canadian Mouse Mutant Repository (The Hospital for Sick Children, 555 University Avenue, Toronto, ON M5G lXB CANADA), and were described before (Hosseini et al, 2022). L7-cre mice were described before (Rico et al, 2004) and were a kind gift of B. Rico (King’s College London, UK). *Ccp1^-/-^* mice were described before (Muñoz-Castañeda et al, 2018).

### Genotyping

Tissue samples for genotyping were small ear biopsies obtained during identification. Tissue was lysed in the lysis buffer (0.1 M Tris–HCl pH 8.0 (Sigma-Aldrich #T1503), 0.2 M NaCl (Sigma-Aldrich #S3014), 5 mM EDTA (Euromedex #EU0007-C) and 0.4% SDS (Euromedex #EU0660) containing 0.1 mg/ml proteinase K (MP Biomedicals #193504) at 65°C for 4 h, followed by proteinase K inactivation (95°C for 10 min). Genotyping was then performed by PCR: for each 10 μl PCR reaction, the mixture contained 5 μl of Platinum polymerase mix (ThermoFisher #14001013), 0.2 μl of DNA extract, 0.06 μl of a premixed forward and reverse primer pair (100 μM), and 5 μl ultrapure H₂O. Positive controls representing all expected genotypes were included in each run. PCR amplification was performed using the following conditions: denaturation: 94°C, 15 s; [annealing: 60°C, 15 s; elongation: 68°C, 40 s] × 30 cycles. Primer sequences and corresponding product sizes are listed in Table 1. Amplified products were separated on a 2% agarose gel containing ethidium bromide and visualized under UV illumination.

**Table 1:**
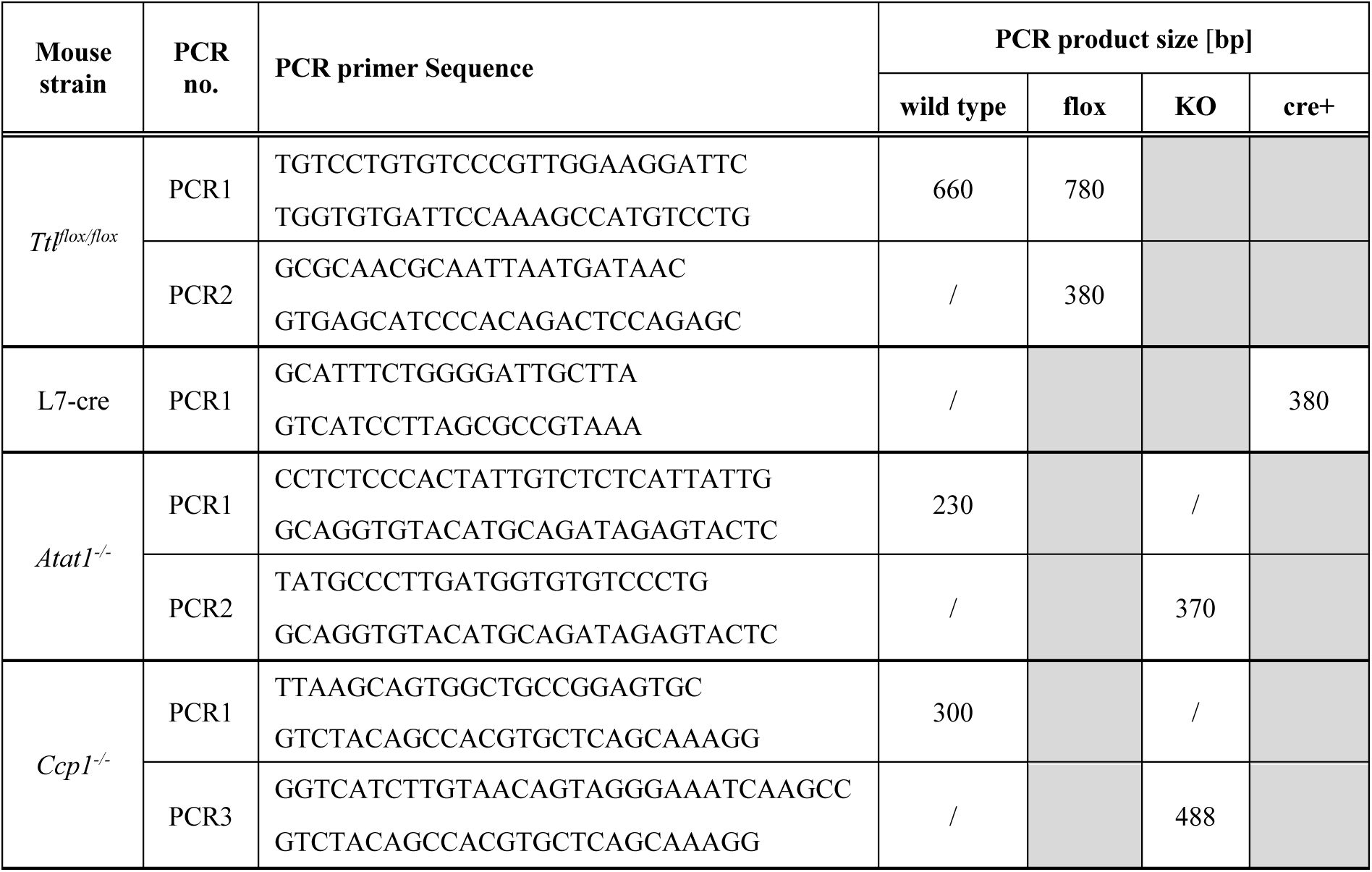
Genotyping primer sequences and expected PCR products sizes.

### Transcardial perfusion

Animals were subjected to transcardial perfusion under deep anaesthesia. Mice were anesthetized with an intraperitoneal injection of a ketamine (Imalgène^®^, 80–100 mg/ml) / xylazine (Rompun^®^, 5–10 mg/ml) mixture (100 µL per 10 g of body weight). Depth of anaesthesia was verified by the absence of the paw withdrawal reflex. The thoracic cavity was opened to expose the heart, after which a needle was carefully inserted into the left ventricle, and a small incision was made in the right atrium to allow venous outflow. Perfusion (at 2-3 ml/min) was initiated with 25-30 ml of 1× PBS until the liver appeared pale and the efflux fluid was clear. Without interrupting the flow, animals were then perfused with 100 ml of fixative solution: freshly prepared 4% paraformaldehyde (Euromedex #15174) in 1× PBS. Following perfusion, brains were rapidly dissected and post-fixed in 10% neutral buffered formalin (Qpath #FOR0020AF59001) at room temperature for 48 h. Subsequently, tissues were washed three times in 1× PBS (30 min per wash) to remove residual fixative and stored at 4°C until further histological or immunohistochemical processing.

### Tissue embedding, immunofluorescence on paraffin sections and imaging

Following post-fixation, brain tissues were washed in 70%, 80% and 95% ethanol for 3 h each, and incubated overnight in 100% ethanol (VWR #64-17-5). Samples were then incubated in isopropanol (Honeywell #33539) for 3 h and infiltrated with molten paraffin wax. Half of the brain was embedded in fresh paraffin using metal moulds, with careful adjustment of tissue orientation using pre-warmed forceps. The moulds were cooled on a cold plate (4°C) for at least 30 min to ensure complete solidification of the paraffin blocks and embedded in cassettes (CellPath #EAI-0611-10A).

Paraffin-embedded brains were sectioned using a microtome (Leica, #RM2245) and 10 µm sagittal sections were collected on glass slides. They were then deparaffinised, rehydrated, and rinsed in TBS-T (Tris-buffer saline with 0.1% Triton X-100). Antigen retrieval was performed by submerging slides in sodium citrate buffer (citric acid 0.1 M (Sigma-Aldrich #251275), sodium citrate tribasic dihydrate 0.1 M (Sigma-Aldrich #S4641), pH 6.0) and microwaving at 800 W for 4 min, followed by 20 min at 145 W. Slides were cooled on ice for 30 min and rinsed in TBS-T, then incubated in the blocking solution (2% BSA, 10% NGS, 0.3% Triton X-100 in TBS) for 2 h at room temperature.

Primary antibodies (Table 2) were applied overnight at 4°C in blocking solution. After washing, secondary antibodies (Table 2), along with 1 µg/ml DAPI were applied for 2 h at room temperature. Slides were mounted using ProLong™ Gold Antifade (Invitrogen #P36934) and imaged on the Nipkow spinning-disk confocal system (Yokogawa CSU-W1) mounted on a Nikon Eclipse Ti2 inverted microscope. Imaging was performed using either a 5× Plan (NA 0.12), 20× Plan Fluor dry objective (NA 0.75), or a 100× Plan Apochromat oil-immersion objective (NA 1.45). Images were captured with a Prime BSI sCMOS camera (Photometrics; pixel size: 6.5 µm) and controlled using MetaMorph® software (Molecular Devices).

**Table 2:** Antibodies used in this study.

| **Antibody** | **Antigen** | **Dilution** | **Reference** |
| --- | --- | --- | --- |
| anti-calbindin (CB-955)<br>mouse monoclonal IgG1 | bovine kidney calbindin-D | 1/200 | Sigma-Aldrich #C9848 |
| anti-calbindin<br>rabbit polyclonal | calbindin CB38a | 1/200 | Swant #AB_3107026 |
| anti-tyrosinated tubulin<br>(TUB-1A2)<br>mouse monoclonal IgG3 | carboxy-terminal amino acids<br>of α-tubulin | 1/200 | Sigma-Aldrich<br>#T9028 |
| polyglutamylation<br>polyE<br>rabbit polyclonal | 3 or more C-terminal<br>glutamates (Rogowski et al,<br>2010), | 1/200 | AdipoGen<br>#AG-25B-0030 |
| Goat anti-rabbit IgG (H+L)<br>Cross-Adsorbed,<br>Alexa Fluor 488 | rabbit IgG | 1/500 | Molecular probes<br>#A11008 |
| Donkey anti-Mouse IgG<br>(H+L) Highly Cross-Adsorbed, Alexa Fluor™<br>647 | mouse IgG | 1/500 | Molecular probes<br>#A31571 |

### Quantification of Purkinje cells

Purkinje cell densities were quantified on 20× magnification images of cerebellar folia IX and X. The length of the Purkinje cell layer in each folium was measured using the segmented line tool in ImageJ (NIH) and the number of Purkinje cells located along the line was manually counted. Purkinje cell density was then calculated as the number of Purkinje cells per mm. Graphic representation and statistical analysis was performed using GraphPad Prism 10, using the unpaired t test with statistical significance set at p<0.05.

### Nissl staining and imaging

Nissl staining was performed on 10 µm paraffin sections which were incubated in 0.1% cresyl violet acetate (Sigma-Aldrich #C5042) for 15 min at room temperature. Excess dye was removed by rinsing in distilled water, followed by differentiation in 70% ethanol for 3 min. Dehydration was performed in ascending ethanol (70% for 3 min, 80% for 1.5 min, 95% for 1.5 min, 100%, for 1.5 min), and sections were cleared in xylene for 4 min. Slides were mounted in the Entellan mounting media (Sigma-Aldrich #1079600500) and imaged using the brightfield Zeiss Axio Imager microscope. Images were obtained at 5×, and 63× magnifications to visualise neuronal morphology.

### Rotarod test

The rotarod test was performed using the 6-cm diameter rod. On day 1, animals were habituated to the room in which the experiment is performed, on day 2 they were trained, and the test itself was performed on day 3. During the training phase, animals were submitted to 4 sessions spaced by at least one hour. During session 1 and 2 they were put on the rotarod rotating at 4 rpm during 3 min, after which the rotation speed was accelerated until 10 rpm over 4 min, and kept at 10 rpm for 4 more min. During sessions 3 and 4 the maximum speed was 15 rpm. The sessions were terminated either after 8 min, or when the animal refused to continue. During the actual test on day 3, animals were put on the rotarod rotating at 4 rpm during 1 min, then the rotation was accelerated progressively to 20 rpm over 3 min. The test was repeated three times for each animal. The time each animal spent on the rotating rod was measured in each test, and the mean of the three tests was calculated. Medians from all mice were plotted and mean values with SEM, as well as statistical analyses were performed with GraphPad Prism 10, using the unpaired t test with statistical significance set at p<0.05. As the motor performance of *pcd*/*Ccp1^-/-^*mice has been extensively tested by rotarod tests in multiple studies analysed in Fig. 3C, we refrained from repeating these experiments to limit the use of animals with damaging phenotypes.

**Supplementary Movie 1. *Ttl^flox/flox^* L7-cre mice do not exhibit ataxic phenotypes.** Representative movies of free-moving control (*Ttl^flox/flox^*), *Ttl^flox/flox^* L7-cre and *Ccp1^-/-^* mice. Animals were put in a clean cage and were filmed exploring the new environment. Note the ataxic phenotype of the *Ccp1^-/-^* mouse, and the normal behaviour of the *Ttl^flox/flox^* L7-cre animals.

